# Sorghum Metabolic Atlas: Large-Scale Subcellular Localization Resource for Sorghum Metabolic Enzymes

**DOI:** 10.1101/2025.08.24.672047

**Authors:** Purva Karia, William P Dwyer, Ava Kloss-Schmidt, Charles Hawkins, Bo Xue, Daniel Ginzburg, Maxine L Gutierrez, Ritesh Mewalal, Ian Blaby, David W Ehrhardt, Seung Y Rhee

## Abstract

Plant metabolism drives traits essential for productivity and resilience, yet understanding metabolic networks requires subcellular, cellular, and tissue-level spatial context that remains limited, particularly in crop species. Experimentally-derived subcellular localization data for enzymes are sparse, constraining analyses of metabolic organization in the cell. We developed a high-throughput protoplast transformation and fluorescent protein (FP) tagging system optimized for *Sorghum bicolor*, a climate-resilient C_4_ crop. Using this platform, we experimentally determined the subcellular localization of 234 metabolic enzymes spanning 184 pathways. The sorghum enzymes we characterized localize to 12 subcellular compartments. Comparison with computational predictions highlights variable accuracy across compartments, and cross-species comparison with *Arabidopsis thaliana* shows partial agreement with available experimental data. All data are accessible through the Sorghum Metabolic Atlas (www.sorghummetabolicatlas.org) web platform, enabling search, visualization, and download. This study presents a large-scale experimental dataset of enzyme localization in sorghum, providing a resource for studies of plant metabolic organization and comparative analyses.

## Introduction

The ability to engineer plant metabolism is essential for adapting agriculture to the challenges posed by rapid climate change. As the primary source for the world’s calories and essential macronutrients, fiber, pharmaceuticals, and biofuel, plants play a critical role in global economies and food security (Mahapatra et al., 2021; Ulian et al., 2020). However, as weather patterns shift and water resources become more scarce, crop yields are predicted to decline significantly, threatening global food supplies (Wing et al., 2021). Synthetic biology and bioengineering offer promising solutions to develop crop varieties with enhanced resilience to these challenges. For example, targeted genome editing via CRISPR-Cas9 has already been used to increase grain yield in rice (Lacchini et al., 2020), improve crop abiotic stress tolerance in tomato (Bouzroud et al., 2020), maize (Shi et al., 2017), rice and wheat (Nazir et al., 2022), and optimize photosynthetic efficiency in rice (Matsumura et al., 2020). Advances in our ability to engineer metabolic pathways are also allowing us to produce useful and novel molecules in plants. For example, engineered biosynthetic pathways have successfully produced medicinal compounds in a heterologous host (Lau & Sattely, 2015; Nett et al., 2020). Despite these advances, a substantial fraction of plant metabolism remains uncharacterized, limiting the scope of targeted metabolic engineering. A broader understanding of the cellular organization of plant metabolism is therefore vital to accelerate these efforts.

A critical function of cellular organization is the compartmentalization of metabolic processes, which enables cells to optimize reaction rates, regulate substrate availability, and mitigate the accumulation of cytotoxic intermediates. By spatially organizing biochemical reactions, at both subcellular and multicellular scales, compartmentalization enhances both efficiency and adaptability of metabolic networks (Emenecker et al., 2020). In C4 photosynthetic plants, for example, the division of photosynthetic reactions between two distinct cell types increases water use efficiency and drought tolerance (Sage, 2004). Pathway flux can be increased by the formation of metabolons, transient complexes of enzymes of a pathway, which are assembled through scaffolding proteins (Boccia et al., 2024). Inspired by this phenomenon, synthetic approaches have been developed to mimic such spatial organization, where enzymes within a metabolic pathway are tethered to an engineered scaffold. This strategy has been shown to significantly enhance pathway flux and product accumulation *in vitro (Geraldi et al., 2021)*. Compartmentalization is also crucial for managing toxic intermediates. By sequestering reactive compounds within specific organelles, cells can prevent cellular damage and optimize metabolic efficiency. In yeast (*Saccharomyces cerevisiae*), for instance, targeting norcoclaurine synthase (NCS) to peroxisomes alleviated its cytotoxicity and boosted alkaloid production (Grewal et al., 2021). These examples highlight the importance of spatial organization in metabolism and its relevance to metabolic engineering approaches.

While numerous studies report subcellular localization data for individual proteins, the broader spatial organization of enzymes remains poorly understood, especially in photosynthetic organisms beyond the model organisms *Arabidopsis thaliana* and *Chlamydomonas reinhardtii*. Computational methods can predict localization with some accuracy, but experimental approaches, such as fluorescent protein tagging, provide higher confidence. Extensive subcellular localization data exist for *A. thaliana*, with over 3500 localized proteins, including 1005 metabolic enzymes (C. Hooper et al., 2022; C. M. Hooper et al., 2017). Recent advances in Chlamydomonas research identified the localization of 1034 chloroplast proteins and uncovered novel localization signals within the chloroplasts using fluorescent protein tagging (Wang et al., 2023). In comparison to Arabidopsis and Chlamydomonas, similar localization efforts in crop species remain limited. Subcellular localizations of only 155 *Oryza sativa* (rice) enzymes, 69 *Zea mays* (maize) enzymes, and just 5 *Sorghum bicolor* (sorghum) enzymes are available in public databases (C. M. Hooper et al., 2016). This scarcity of enzyme localization data in crops impedes our ability to engineer plant metabolic pathways and improve crop varieties.

*Sorghum bicolor*, the fifth most-produced cereal crop globally, is an understudied *Poaceae* family member with higher resilience to abiotic stresses such as drought and heat than other cereal crops (Ali et al., 2023; Kim et al., 2019). Currently, only three cereal crop species, wheat, maize, and rice, supply over a third of the world’s calories (Shiferaw et al., 2011). Despite high yields and a long history of cultivation, these crops are resource intensive and predicted to lose productivity in warmer, drier climates (C. Zhao et al., 2017). Expanding the use of stress-tolerant crops into agricultural systems is a promising strategy to buffer food supply chains against climate shocks.

Chief among these crop candidates is sorghum, a naturally drought-and heat-resistant, nitrogen-use efficient cereal crop originating from eastern Africa (Venkateswaran et al., 2019). Sorghum is also a highly versatile crop. In addition to the nutritious grain for which it is cultivated, the plant is also a source of animal feed and a promising candidate in the production of sustainable ethanol-based biofuels (Silva et al., 2022). Despite its natural resilience and promising traits, research on sorghum at the cellular and metabolic level is limited, impeding efforts to fully exploit its potential for agricultural and industrial use.

Given the importance of enzyme subcellular localization and the increasing relevance of sorghum in climate-resilient agriculture, this study aims to address the current gap in localization data for key metabolic enzymes in sorghum. Here, we introduce the Sorghum Metabolic Atlas (SMA), a subcellular map of sorghum enzymes involved in small-molecule metabolism, generated using fluorescent protein (FP) tagging in sorghum protoplasts. This dataset details the localization of 234 enzymes across 12 compartment classes and represents, to our knowledge, the first large-scale effort to localize plant enzymes in sorghum. The SMA provides experimentally supported localization data and establishes a resource for studies of plant metabolic organization and comparative analyses.

## Materials and Methods

### Enzyme selection

Orthology between *Arabidopsis thaliana* and *Sorghum bicolor* proteomes was established using the OrthoFinder pipeline (Emms & Kelly, 2019), with one multi-FASTA file per species. Protein datasets were filtered to retain only metabolic enzymes by using SmartTables within the Plant Metabolic Network database (Hawkins et al., 2021). For each orthogroup containing *A. thaliana* genes, experimental subcellular localization data were retrieved from the Subcellular Localization Database for Arabidopsis Proteins (SUBA) database (C. Hooper et al., 2022). Enzyme candidates were selected to span the breadth of plant metabolism and across three different categories: 1) *S. bicolor* genes with experimentally localized orthologs in *A. thaliana*, 2) *S. bicolor* genes with *A. thaliana* orthologs that lack experimental localization, and 3) *S. bicolor* genes without orthologs in *A. thaliana*. We also reserved a portion of our dataset for pathways of relevance to ongoing research and breeding efforts. When multiple isozymes were present for a given enzymatic reaction, selection was guided by sorghum transcript abundance (Olson et al., 2014).

### Vector assembly

The pGVG (Guidelli et al., 2018) and pEZS (https://deepgreen.dpb.carnegiescience.edu/cell%20imaging%20site%20/html/vectors.html) vectors were obtained from the Menossi and Ehrhardt labs, respectively. These vectors were modified to add a fluorescent tag either upstream or downstream of the coding sequence enabling the expression of the fusion protein from a single transcript. PCR amplification was carried out for three fragments: 1) the vector backbone containing the promoter and terminator, 2) the Gateway cassette, and 3) the fluorescent (mEGFP or mCherry) tag using PrimeSTAR Max DNA Polymerase according to the manufacturer’s instructions. The pGVG vector backbone contains the maize polyubiquitin promoter (Ubi-1) and the Cauliflower mosaic virus 35S promoter (CaMV35S) terminator, while the pEZS vector backbone contains the CaMV35S promoter and the Octapine synthase 3 (OCS3) terminator. PCR fragments were gel-extracted and purified then assembled using the In-Fusion seamless cloning system, following the manufacturer’s instructions (Takara Bio, #638947). Transformants were selected on Luria-Bertani agar plates containing 25µg/mL chloramphenicol for both pGVG and pEZS constructs. The vector assembly was verified by long-read sequencing and the vector maps are shown in Figure S1a-c.

### Cloning and sequence analysis of protein expression constructs

The coding sequences of selected *S. bicolor* enzymes were refactored using an *S. bicolor* codon usage table to overcome synthesis constraints (Oberortner et al., 2017), synthesized (Twist Bioscience, CA) and cloned into the Gateway entry vector pDONR221. To guide fluorescent tag placement, amino acid sequences were analyzed using TargetP 2.0 (Almagro Armenteros et al., 2019) to predict N-terminal targeting peptide for chloroplast, mitochondria, and secretory pathways and Plant PTM Viewer 2.0 (Willems et al., 2024) to identify post-translational protein modifications near protein termini. Based on predicted targeting signals, enzymes with chloroplast transit peptides (cTPs) were C-terminally tagged, while others were N-terminally tagged. For mitochondria and secretory pathway candidates, either N-or C-terminus tagging was opted. A subset of enzymes, primarily those predicted to localize to the chloroplast, were cloned in both N-terminal and C-terminal fusion with the fluorescent tag for localization validation (Figure S1d, Table S1). Coding sequences were designed to maintain the correct reading frame upon recombination with the fluorescent protein tag in the destination vectors. Entry clones were recombined into destination vectors using LR Clonase II (Invitrogen) according to the manufacturer’s instructions. Constructs were sequence-verified at the promoter-coding sequence (CDS) junction using UBIQ-Forward (5’ GGCATATGCAGCAGCTATATGTGGA 3’) or 35S-Forward (5’ CAACCACGTCTTCAAAGCAAGT 3’) primers (for C-terminal fusions) or GFP-Forward (5’ ATGGTCCTGCTGGAGTTCGT 3’) or mCherry-Forward (5’ ATCAAGTTGGACATCACCTCC 3’) primers (for N-terminal fusions), ensuring the correct reading frame, allowing for proper transcription and translation of a continuous fusion protein with the fluorescent tag.

### Protoplast preparation

The protoplast isolation protocol was adopted and modified from (Birnbaum et al., 2005; Gomez-Cano et al., 2019; Shulse et al., 2019). Protoplasts were isolated from 1g of 15-17 youngest fully expanded leaves of 2 to 3-week-old *Sorghum bicolor* RTx430. Leaves were sliced into 1mm-wide strips using a scalpel and transferred to 20mL of Solution A (0.6M mannitol, 2mM MgCl_2_.6H_2_O, 2mM CaCl_2_.2H_2_O, 10mM KCl, 2mM MES hydrate, 0.1% BSA; pH 5.5, adjusted with Tris base) supplemented with 1.25% cellulase R-10 (GoldBio, catalog #C8001.0005) and 0.3% macerozyme R-10 (GoldBio, #M8002.0001). Samples were vacuum-infiltrated twice at 15Hg for 15 minutes each, then incubated in the dark at room temperature for 50 minutes on a horizontal shaker at 100 RPM. To stop cell wall digestion, an equal amount of Solution A was added, the entire mixture was filtered through a 70µm nylon filter (Greiner Bio-One #542070), and centrifuged at 100 x g for 10 minutes at room temperature. At this stage, the supernatant was carefully decanted, cells were washed using 10mL of Solution A, centrifugation was repeated as mentioned above, and the protoplasts were resuspended in 500µl of solution A. Using approximately 1g of leaf tissue in the protocol produced 1.5 - 4.5 x 10^5^ sorghum protoplasts/mL.

### Protoplast yield calculation and viability assessment

Protoplast yield was calculated by measuring protoplast density with a double-chamber hemocytometer (Fisher Scientific #0267110) and the final concentration was adjusted to 10^5^ cells/mL in solution A. Protoplast viability was assessed by staining with 0.01% (w/v) fluorescein diacetate (FDA) for 5 minutes and visualizing under an epifluorescence microscope. Assessment of protoplast viability was performed on 3 independent preparations of protoplasts.

### Plasmid purification and protoplast transformation

Plasmid DNA was purified using the HiPure Plasmid miniprep kit (Invitrogen™, #K210003) according to the manufacturer’s instructions and concentrations were measured by NanoDrop 1000 (ThermoScientific). To introduce plasmid DNA into the protoplasts, PEG-mediated transformation was performed. For PEG-mediated transformation, 50µl of protoplasts, up to 8µL (5µg) of plasmid DNA, and 55µL of 40% PEG transfection buffer (containing 0.6M mannitol, 40% PEG-4000, 0.1M CaCl_2_) were sequentially added to a 1.5mL Eppendorf tube, which was inverted a few times to mix gently. The transformation solution was incubated in the dark at room temperature for 20 minutes. 400µL of solution A was added to the transformation solution and centrifuged at 100 x g for 10 minutes. The supernatant was decanted; cells were washed and resuspended in 1mL of solution A. The cells were then transferred to a 24-well plate for overnight incubation in the dark.

### Sample preparation for microscopy

Transformed protoplasts were transferred to 1.5mL Eppendorf tubes and centrifuged at 110 x g for 10 minutes. Approximately 900-950µl of supernatant was pipetted out. The protoplasts were gently resuspended in the remaining solution. A 12µL aliquot of the suspended protoplasts was added to each well of the CultureWell™ multiwell chambered coverslips (Thermo Fisher, #C24779) and covered with a plain coverslip (VWR, #48366-277). For mitochondrial staining, 0.5µL of 1mM MitoSpy™Red CMXRos (BioLegend) stock was added to 1mL of Solution A. Lipid droplets were stained using either Nile Red or BODIPY. For Nile Red staining, 2µL of 1mM Nile Red (Invitrogen™) stock was added to 1mL of Solution A. For BODIPY staining, 5µL of 1mM BODIPY (Invitrogen™) stock was added to 445µL of protoplast Solution A. For each stain, equal volumes of staining solution and protoplasts were mixed and incubated for 20 minutes before confocal imaging.

### Image acquisition

Image acquisition of transfected protoplasts was performed using a Leica SP8 confocal laser scanning microscope with a glycerine immersion objective and white light laser. Excitation energy and emitted fluorescence were collected using the following parameters: mEGFP - 489nm excitation, 500-550nm emission; mCherry - 587nm excitation, 600-650nm emission; mCitrine - 513nm excitation, 525-570nm emission; chloroplast autofluorescence - 680-700nm emission. To minimize chloroplast autofluorescence, we applied emission time gating (0.7-9.0ns), which excludes short-lived autofluorescence (e.g., from chlorophyll) while capturing the longer-lived fluorescence of mEGFP and mCherry channels. Compatible fluorophore combinations were used for colocalization experiments, and sequential scanning was employed to prevent signal bleed-through among channels.

### Image analysis

Confocal microscopy images were analyzed using Fiji (Schindelin et al., 2012). For each enzyme, a single cell or a small group of cells displaying the dominant localization pattern was selected for analysis. In cases where localization signals were ambiguous or not clearly associated with a distinguishable organelle, colocalization experiments were performed to verify subcellular targeting. Colocalization was quantified using JaCOP (Just Another Colocalization Plugin) in Fiji and evaluated using Manders’ overlap coefficient with Costes automatic thresholding (X. Zhao et al., 2022).

### Web application

The Sorghum Metabolic Atlas website (https://sorghummetabolicatlas.org/) is primarily written in TypeScript 4.4.3 (https://www.typescriptlang.org/docs/handbook/intro.html) and is executed within the <u>Node.js</u> 23.11.1 (https://nodejs.org/en/) runtime environment. The backend uses PostgreSQL 16.9 (https://www.postgresql.org/docs/current/) for data storage and retrieval. The front end uses Angular 13.0.1 (https://angular.dev/), with the front-page cell diagram implemented as SVG drawn using Inkscape 1.2 (https://inkscape.org/). Data updates are handled using bespoke scripts.

## Results

### Selection of enzymes across metabolism for in planta localization

To map metabolism across diverse subcellular compartments in sorghum leaf cells, we selected 417 out of 9443 enzymes from SorghumBicolorCyc version 7.1.1 (Hawkins et al., 2021) based on their metabolic pathway annotation. When multiple enzymes were associated for a given reaction, the isozyme with the highest median expression across all tissues was selected (Olson et al., 2014) (Table S2). Coding sequences were synthesized, cloned into a GFP translational fusion expression vector, transformed into sorghum leaf protoplasts, and imaged using confocal microscopy. Of the 417 candidates, 234 enzymes yielded detectable and assignable localization signals, an efficiency of ∼56%. These represent 184 metabolic pathways, spanning 13 metabolic domains as defined by (Schläpfer et al., 2017) (Table S3). The most represented were the amino acid, energy, and specialized metabolism domains (Figure 1a). This dataset includes enzymes from diverse metabolic pathways across multiple domains and was generated using a consistent experimental pipeline for subcellular localization analysis.

**Figure 1:**
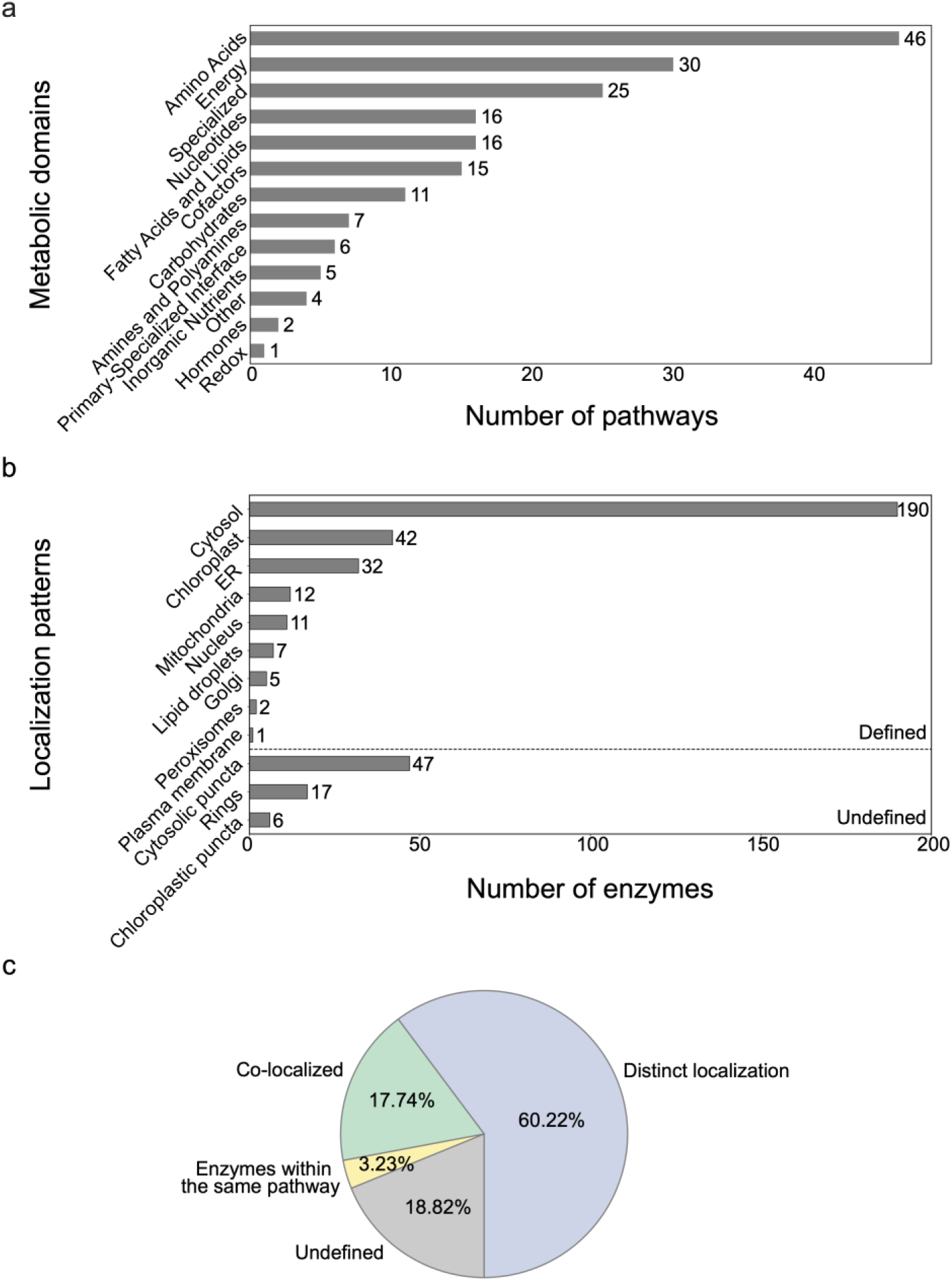
Metabolic classification and subcellular localization of selected *Sorghum bicolor* enzymes. a) Number of pathways represented across major metabolic domains. b) Subcellular localization of enzymes based on fluorescence-tagging sorghum protoplasts. “Defined” and “undefined” localizations are separated by the dotted line. c) Enzyme localization compartments are assigned based on 1) resemblance to known compartment makers as distinct localization, 2) co-localization with compartment markers, 3) pathway context for enzymes within the same pathway, and undefined cases lacking experimental evidence.

### A broad cellular view of sorghum metabolic enzymes

We constructed a subcellular localization dataset of sorghum metabolic enzymes by expressing 234 fluorescently tagged protein-coding sequences in sorghum protoplasts. All enzymes were localized to at least one of 12 subcellular compartment classes (Figure 1b, Table S1): cytosol, cytosolic puncta, chloroplast, chloroplastic puncta, endoplasmic reticulum (ER), rings, nucleus, Golgi apparatus, lipid droplets, mitochondria, peroxisomes, plasma membrane (Figure 1b). Compartment classes that were defined by morphology only, such as cytosolic puncta, chloroplastic puncta, and rings, may represent distinct but uncharacterized functional compartments. About half of the enzymes displayed signals in the cytosol (51%), followed by the chloroplast (11%), ER (9%), mitochondria (3%), and lipid droplets (2%). Smaller subsets were observed in the Golgi apparatus, nucleus, plasma membrane, and peroxisomes. Additionally, some localized enzymes displayed cytosolic puncta (13%), rings (5%), and chloroplastic puncta (2%) with unclear organellar identity (Figure 1b).

Compartmental assignments were supported by three complementary strategies (Figure 1c). Localizations of most enzymes (60%) were classified based on their resemblance to established compartmental markers with distinctive and diagnostic distributions, such as cytosol and ER (Figure S2a). Another 18% were supported by direct co-localization with fluorescent compartmental markers (Figure 1c, Figure S2b). 3% of enzyme localizations were inferred from metabolic pathway context, where at least one enzyme co-localized with a compartment-specific marker and other pathway members displayed consistent fluorescence patterns (Figure 1c). For example, several enzymes involved in leucine degradation exhibited similar localization signals, supporting their assignment to mitochondria (Figure S2c-e). The remaining 19% displayed patterns that were not easily assigned to known organelles, including cytosolic puncta, rings, and chloroplastic puncta (Figure 1c, Figure S2f-g). Assignment strategies were tailored to each compartment, with localization annotation summarized by both the compartment and classification method (Figure S2h). Overall, these methods and this dataset provide a consistent experimental framework for assigning enzyme localization in sorghum, while identifying candidates with unresolved or complex localization for future study.

### Multi-compartment localization of metabolic enzymes

A notable finding from our dataset is the widespread occurrence of multi-compartment localization among sorghum metabolic enzymes. We analyzed 254 localization observations derived from N-or C-terminally tagged constructs representing 234 unique enzymes. Of these, 153 (60%) localized to a single compartment, while 101 (39%) localized to multiple compartments including 85 (33%) with dual and 16 (6%) with three or more compartments (Figure 2a). We observed two distinct classes of multi-compartmental localization: 1) signal in multiple compartments within the same cell, and 2) signal in distinct compartments in different cells. It is possible that multi-compartment localization could arise from the overexpression of tagged proteins. Multi-compartment localization may reflect dynamic or conditional subcellular distribution, potentially influenced by differences in cell types or physiological states. However, we cannot exclude the possibility that overexpression of fluorescently tagged proteins may saturate the native targeting machinery, leading to partial mislocalization. Likewise, puncta might form a localization class when proteins are overexpressed due to misfolding and aggregation.

**Figure 2:**
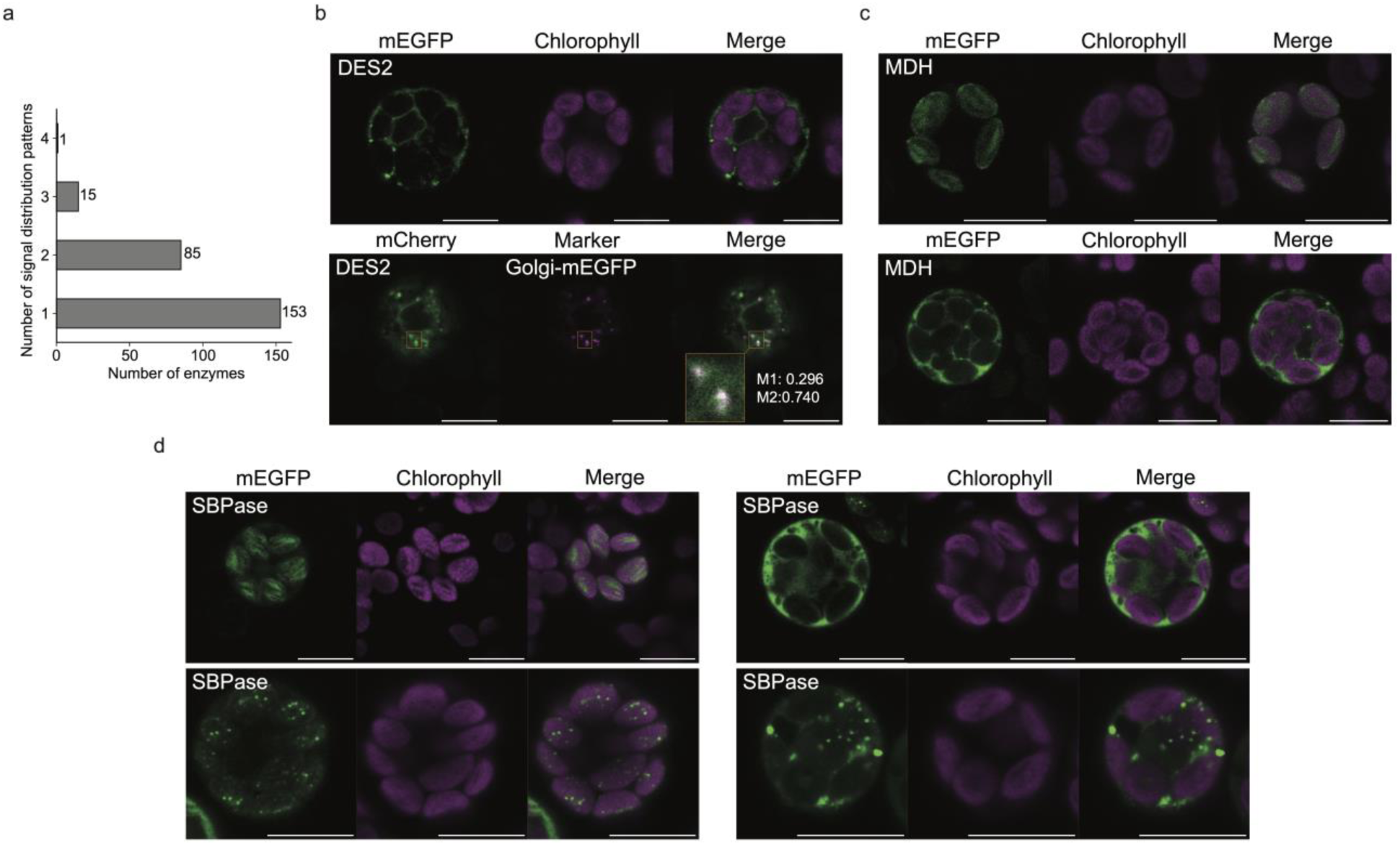
Subcellular localization patterns of enzymes across multiple compartments. a) The bar graph represents the number of enzymes associated with one or multiple localization patterns. b) Sphingolipid 4-monooxygenase (DES2-Sobic.002G383900) is located in two compartments within the same cell. Top: Confocal images showing fluorescence signals of the fluorescent protein (FP) (green) in ER and Golgi (puncta). Bottom: Co-localization with a Golgi marker further confirms its localization to the Golgi apparatus. c) Malate dehydrogenase (MDH-Sobic.007G166300) is located in different compartments in different cells. Representative images showing localization in distinct subcellular compartments across different cells, as indicated by mEGFP fluorescence (green) and chlorophyll autofluorescence (magenta). d) Sedoheptulose bisphosphatase (SBPase-Sobic.003G359100) is located in four distinct compartments. Confocal images showing fluorescence signals for FP (green), chlorophyll autofluorescence (magenta), and their merge. The protein localizes to the chloroplast, chloroplast-associated puncta, cytosol, and puncta within the cytosol. Scale bars represent 10 µm.

Examples from our dataset highlight both technical and biological sources of localization diversity. Sphingolipid 4-monooxygenase (DES2; Sobic.002G383900), an enzyme that introduces double bonds into sphingolipid backbones, was found in both ER and Golgi-associated puncta. Co-localization using a Golgi marker confirmed its presence in the Golgi (Figure 2b), consistent with the enzyme’s known role in sphingolipid biosynthesis, which spans the ER-Golgi interface. However, we cannot exclude the possibility that the ER localization could also result from overexpression-induced retention in the ER lumen. In another example, malate dehydrogenase (MDH; Sobic.007G166300) catalyzes the formation of malate, the major transport metabolite of NADP-ME type C_4_ photosynthesis. Malate carries fixed carbon from mesophyll chloroplasts to bundle sheath chloroplasts, where decarboxylation releases CO_2_ for Rubisco, making MDH an important component of the C_4_ carbon-concentrating mechanism. When C-terminally tagged, MDH was found either in the cytosol or chloroplast in different cells (Figure 2c). The cell-to-cell variation may arise from differential protein import efficiency or reflect true differences in localization across distinct cell types.

More complex multi-compartmental localization was evident for sedoheptulose bisphosphatase (SBPase; Sobic.003G359100), a key Calvin-Benson cycle enzyme involved in regenerating ribulose-1,5-bisphosphate in bundle sheath cells. Signals were detected in up to four compartments, most prominently in the chloroplast and cytosol, but also as diffuse or puncta patterns that differed from cell to cell (Figure 2d). These observations may reflect potential compartment-specific functions in regulating the enzyme among different cell types or states. However, we cannot rule out the possibility that variable formation of puncta could be caused by overexpression. These observations highlight both the subcellular spatial complexity of enzyme localization in plant cells, and the need to develop methods for in-planta localization using endogenous tagging.

### Assessment of computational localization prediction and cross-species comparison

An important goal of the Sorghum Metabolic Atlas is to provide experimentally supported localization data for benchmarking and improving computational prediction tools, including those generated by artificial intelligence (AI). To assess the accuracy of localization predictions, our experimentally determined localizations were compared with TargetP predictions for chloroplasts, mitochondria, and the secretory pathway, compartments that are routinely predicted by algorithms. Prediction accuracy varied by compartment, with true positive rates of 66% for chloroplast, 53% for the secretory pathway, and 25% for mitochondria (Figure 3a). Additionally, a subset of enzymes was experimentally observed in these compartments despite being predicted elsewhere, indicating false negatives. These findings highlight the utility of computational tools while emphasizing the need for experimental validation, particularly for mitochondrial targeting.

**Figure 3:**
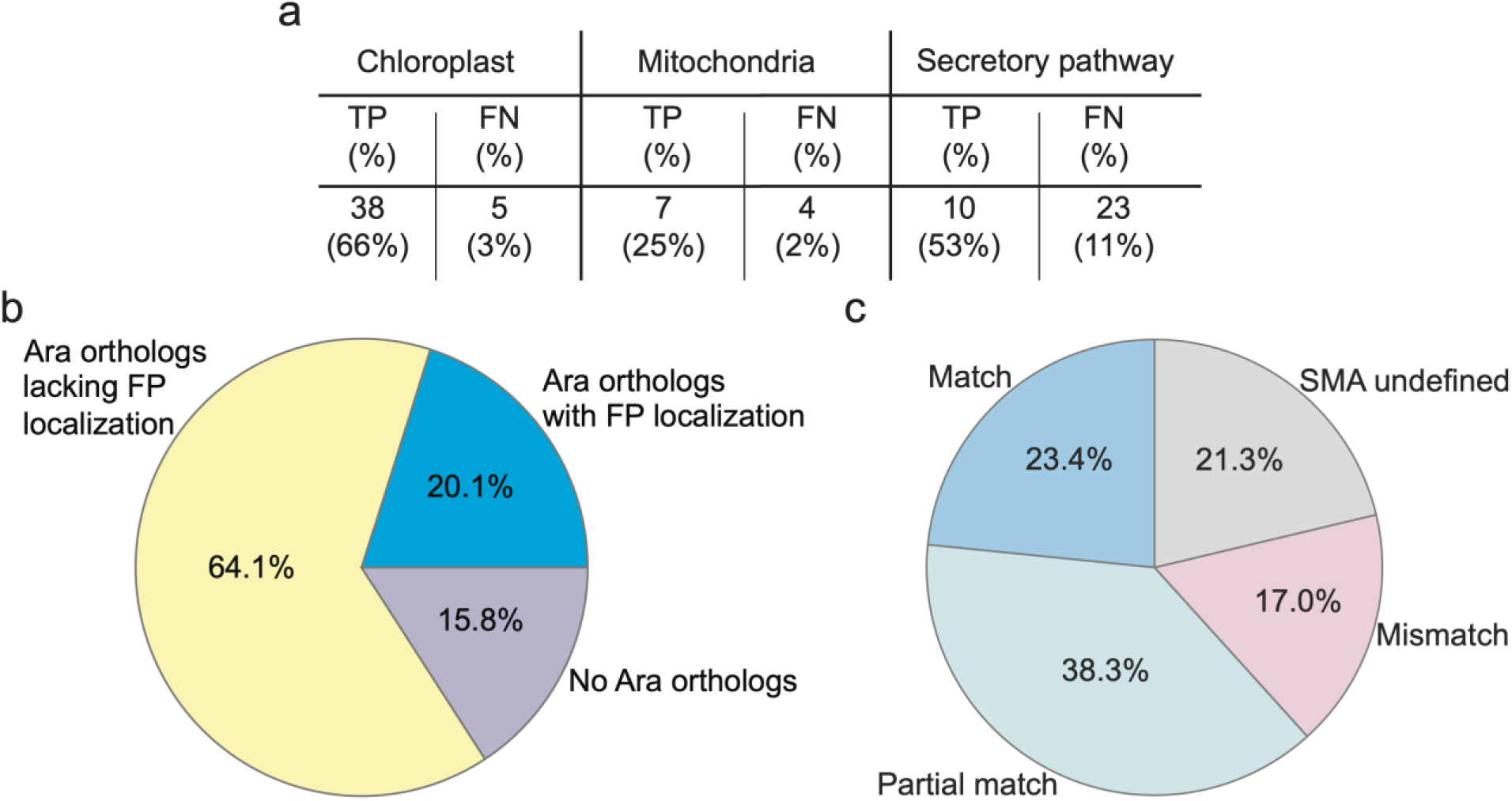
Localization accuracy and cross-species conservation between sorghum and Arabidopsis. a) Comparison of experimentally determined enzyme localization with TargetP predictions for three subcellular compartments: chloroplast, mitochondria, and secretory pathway. True positives (TP) indicate agreement between prediction and experimental localization, while False negatives (FN) indicate predicted localization not matching observed localization. Percentage reflects the proportion of the total enzyme dataset. b) Classification of enzymes based on orthology with Arabidopsis genes and availability of fluorescence protein (FP) localization data: Arabidopsis orthologs with FP localization data, Arabidopsis orthologs lacking FP localization data, and enzymes with no identifiable Arabidopsis orthologs. c) Comparison of sorghum enzyme localization with the localization of Arabidopsis orthologs grouped as: Match - same compartment in both species; Partial match - localization to multiple compartments in one or both species with only partial overlap; Mismatch - localize to different compartments; SMA unidentified - localization data from Sorghum Metabolic Atlas (SMA) ambiguous, preventing direct comparison with Arabidopsis.

To assess our localization data relative to the existing enzyme subcellular location knowledge, we compared the sorghum enzyme dataset with *A. thaliana* enzymes that have been localized using fluorescent protein (FP)-based localization. Of the 234 enzymes, 197 (84%) have orthologs in Arabidopsis. Among the 197 Arabidopsis orthologs, only 47 (24%) have FP-based localization data in SUBA (Figure 3b). Most of the sorghum enzymes localized to the same compartments as their Arabidopsis homologs, with 24% matching exactly, 38% matching partially (defined as localization to at least one shared compartment), and 17% showed distinct localizations. The remaining 21% of sorghum enzymes lacked directly comparable or well-defined localization patterns and were excluded from this comparison. The proportion of matching and partially matching localizations suggests consistency with previously reported localization data, extent of enzyme location conservation, and reliability of our experimental pipeline. Overall, our dataset substantially extends available localization information in a crop species and helps address gaps in experimentally validated subcellular localization data in plants.

### Data accessibility through the Sorghum Metabolic Atlas (SMA) database

To facilitate dissemination of these data and enable hypothesis generation by the research community, a searchable Sorghum Metabolic Atlas (SMA) web application has been created. Publicly available at sorghummetabolicatlas.org, the SMA website allows users to search for sorghum proteins and find their subcellular localization that is supported by fluorescent protein tagging. The SMA enables users to search sorghum enzymes and view experimentally determined subcellular localizations supported by fluorescent protein tagging, or to search by organelle to identify localized proteins. The database contains 297 images representing localization data for 234 sorghum enzymes and includes additional information such as amino acid sequences, associated reactions, and pathway annotations from SorghumBicolorCyc metabolic pathway database (https://pmn.plantcyc.org/organism-summary?object=SORGHUMBICOLOR), which is part of the Plant Metabolic Network (Hawkins et al., 2021). Protein entries can be searched and filtered by criteria including protein ID, subcellular localization, Arabidopsis homologs, metabolic pathways, and catalyzed reactions, and filtered datasets may be exported to a comma-separated values (CSV) file for further analyses.

## Discussion

Using a high-throughput fluorescent protein (FP) tagging approach in sorghum protoplasts, we generated the Sorghum Metabolic Atlas (SMA), an expansive dataset capturing localization information for 234 enzymes across 184 pathways and 12 distinct subcellular compartments. This resource captures a range of localization patterns that provide data for investigating metabolic organization and regulation at the cellular level. The online SMA database (sorghummetabolicatlas.org) enables the broader plant biology community to access these data in a user-friendly way. The image and annotation data may be useful for benchmarking and improving localization prediction algorithms. This study provides a dataset for examining the subcellular organization of metabolism in *Sorghum bicolor*, provides a dataset relevant to studies of plant metabolism and metabolic engineering in sorghum and other C_4_ crops.

Large-scale protein localization efforts have been conducted in model systems such as *A. thaliana (Cutler et al., 2000; Tanz et al., 2013)*, *C. reinhardtii (Wang et al., 2023)*, yeast (Huh et al., 2003), and human cell lines (Thul et al., 2017). However, comparable datasets for crop species are severely limited, with maize being a notable exception (Krishnakumar et al., 2015). This gap has limited our understanding of the cellular organization of plant metabolism and its variation across plant lineages. By focusing on *Sorghum bicolor*, an important bioenergy crop, this study addresses a critical gap in plant systems biology and offers a reference point for comparative studies across species.

To enable subcellular localization in sorghum, we established a protoplast-based pipeline that enabled the generation of localization data across diverse metabolic pathways. However, limitations such as the absence of cell walls, use of constitutive promoters, and potential steric effects from FP tags may influence observed localization patterns. Of 417 enzymes selected, 234 yielded detectable and assignable localization signals, while 183 failed to produce interpretable localization patterns, often due to weak fluorescence, low protein accumulation, or other technical limitations associated with transient expression and imaging. Over 42% of the enzymes exhibited multi-compartment localization (Figure 2a), which may reflect biological complexity or artifacts from overexpression, tagging, or misfolding. For instance, a single isoform of malate dehydrogenase (MDH) localized to either the cytosol or chloroplast in different cells (Figure 2c). This variability may reflect differences in protein targeting efficiency, cell type, or cell state. Further investigation, including the use of cell-type-specific markers, will be required to resolve these patterns in the future.

The SMA is designed as a community resource to support hypothesis generation, comparative analyses, and data integration. The availability of fluorescent images, localization annotations, and associated metadata in a searchable format enables exploration of enzyme localization across pathways and compartments, and integration of these data into broader studies of plant metabolism. Future extension of this resource could include integration with additional cell types, use of stable expression systems, and comparison to other species.

Overall, this study highlights both the capabilities and constraints of FP-based approaches for characterizing enzyme localization. By providing experimentally-derived localization data in a crop species, the SMA supports studies of plant metabolic organization and evaluation of computational prediction methods. This resource establishes a foundation for further investigation of enzyme localization and its role in plant metabolism. As climate change intensifies the demand for resilient, resource-efficient crops, spatially resolved metabolic information may contribute to future efforts in plant engineering.

## Acknowledgements

This work was funded largely by the U.S. Department of Energy, Office of Science, Office of Biological and Environmental Research, Genomic Science Program grant DE-SC0020366 to SYR and DWE. Also the work was supported in part by the U.S. National Science Foundation grants (IOS-2312181, IOS-2406533, IOS-1546838, MCB-1617020, MCB-2052590, MCB-1916797, MCB-2420360, OISE-2434687, DBI-2213983 to SYR) and U.S. Department of Energy, Office of Science, Office of Biological and Environmental Research, Genomic Science Program grants (DE-SC0018277, DE-SC0023160, DE-SC0008769, and DE-SC0021286 to SYR). The work (proposal: https://doi.org/10.46936/10.25585/60000007) was conducted by the U.S. Department of Energy Joint Genome Institute (https://ror.org/04xm1d337), a DOE Office of Science User Facility, is supported by the Office of Science of the U.S. Department of Energy operated under Contract No. DE-AC02-05CH11231. We thank the members of the Sorghum Metabolic Network project for their insights and helpful discussions related to this work. We thank members of the Rhee lab and the Plant Cell Atlas community for thoughtful discussion throughout this project. We thank the Ehrhardt lab for providing the pEZS vector backbone, the Menossi lab for the pGVG vector backbone, and the Kodama lab for organelle markers (cytosol, nucleus, and mitochondria). We thank Andrey Malkovskiy at the Carnegie Science Microscopy Core for assistance with microscopy. We acknowledge Maxim Koriabine from the DOE Joint Genome Institute in Berkeley for the valuable discussions and insights regarding the protoplast isolation protocol. This work was conducted in part on the ancestral land of the Muwekma Ohlone Tribe which was and continues to be of great importance to the Ohlone people, and on the ancestral, traditional, and contemporary lands of the Anishinaabeg – Three Fires Confederacy of Ojibwe, Odawa, and Potawatomi peoples. The work was also conducted at Michigan State University that occupies the ancestral, traditional, and contemporary Lands of the Anishinaabeg – the Three Fires Confederacy of Ojibwe, Odawa, and Potawatomi peoples. We affirm Indigenous sovereignty and hold Michigan State University accountable to the needs of American Indian and Indigenous peoples.

## Competing interests

Authors declare no competing interests.

## Author contributions

Conceptualization: PK, SYR, DWE Methodology: PK

Software: BX, CH

Investigation: PK, SYR, AKS, WPD, DG, MLG, BX

Visualization: PK, AKS, WPD, BX, CH Resources: SYR

Funding acquisition: SYR, DWE, WPD Project administration: SYR Supervision: PK, SYR, IB, DWE Writing – original draft: PK, AKS, WPD

Writing – review & editing: PK, SYR, AKS, WPD, IB, DWE, MLG, BX, DG

## Data availability

Plasmids and constructs described in this work will be deposited in Addgene. Enzyme-related metadata and imaging resources are publicly available at sorghummetabolicatlas.org.

## Supporting Figures

**Figure S1:**
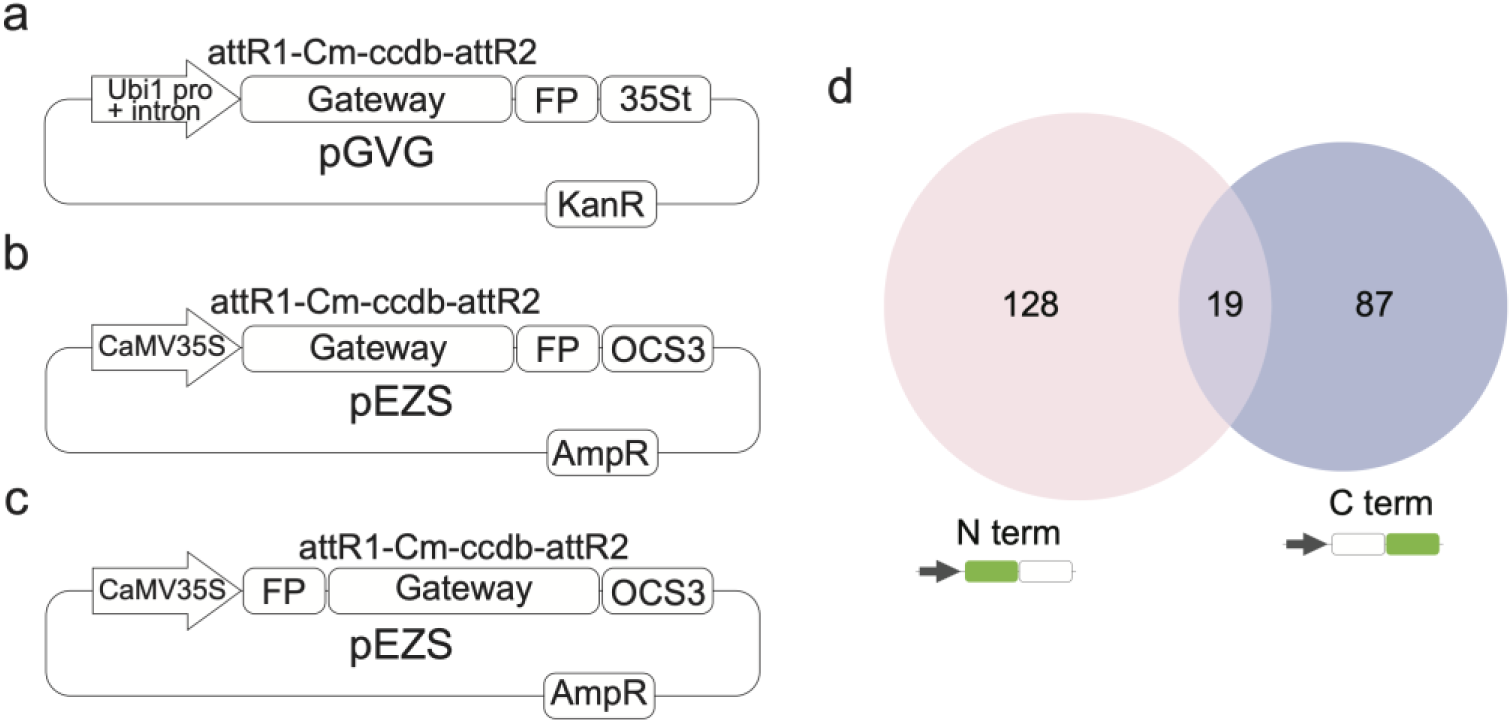
Vector design and comparison of N and C terminal fusion constructs. a-c) Schematic diagrams of Gateway-compatible binary vectors used for subcellular localization in sorghum protoplasts. a) pGVG vector containing the maize Ubiquitin-1 promoter with intron, C-terminal mEGFP tag, and 35S terminator, conferring kanamycin resistance. b,c) pEZS vector containing CaMV 35S promoter, FP fusion at (b) C-terminal or (c) N-terminal, OCS3 terminator, conferring ampicillin resistance. d) Venn diagram showing sorghum enzymes tagged with fluorescent proteins at the N- or C-terminus for localization analysis.

**Figure S2:**
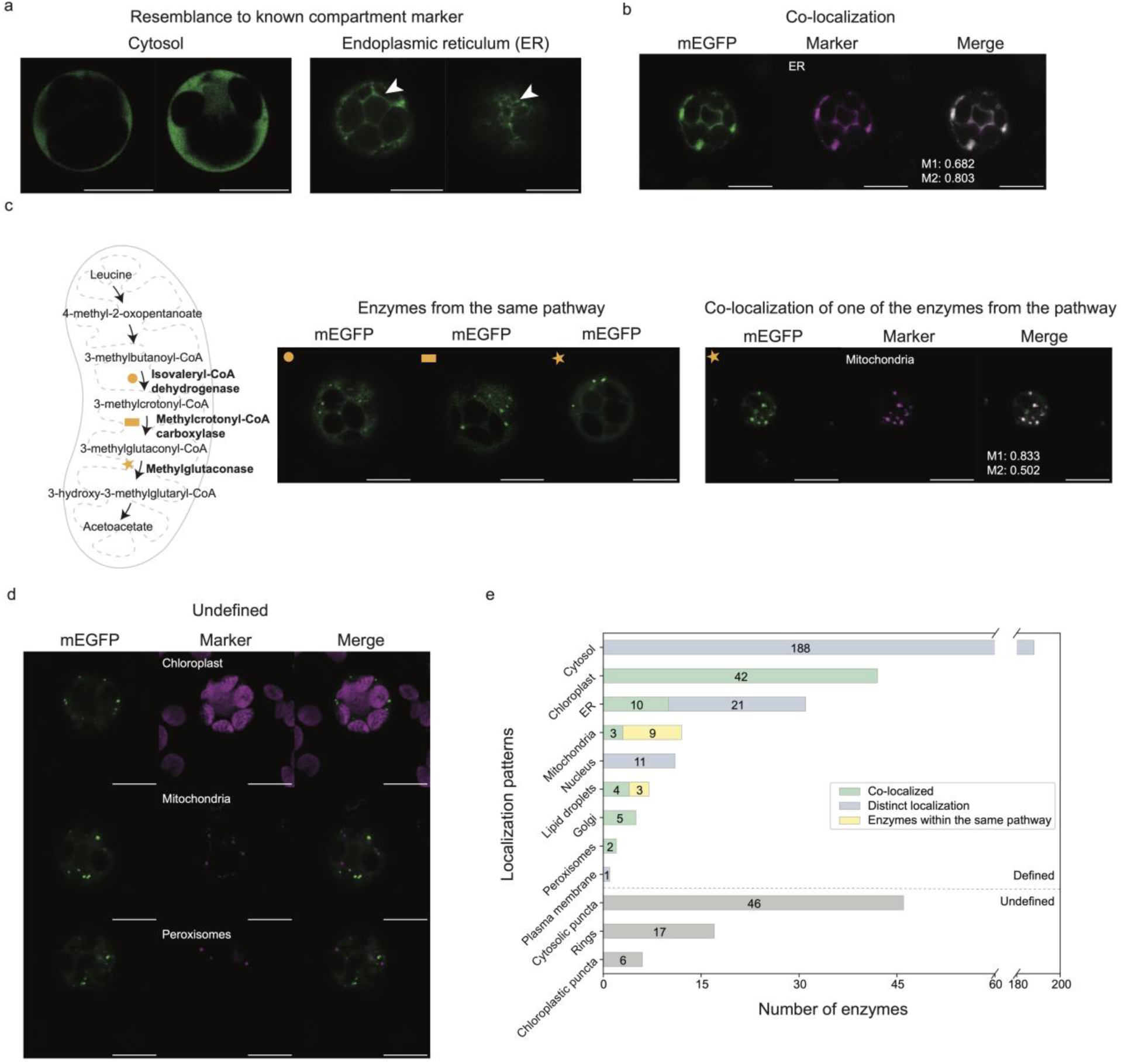
Assigning subcellular compartments based on localization patterns. a) Representative images of patterns of fluorescent-tagged enzymes showing cytosolic (Sobic.008G179900) or endoplasmic reticulum (Sobic.004G064000). 2 sections from the z-stack are shown for each enzyme. b) Representative image of co-localization of GFP-tagged enzyme (Sobic.001G195100) with an ER-marker, confirming ER localization. c) Schematic of leucine degradation pathway with analyzed enzymes highlighted (in bold) include Sobic.009G027000, Sobic.007G130600, and Sobic.004G297400. d-e) Highlighted enzymes show puncta and weaker cytosolic signals. Methylcrotonyl-CoA carboxylase co-localizes with the mitochondrial marker, confirming mitochondrial localization. f) Enzymes with undefined cytosolic puncta and rings are shown via localization patterns of Cysteine Desulfurase (CD-Sobic.004G132600) and Succinate Semialdehyde Dehydrogenase (SSADH-Sobic.004G058600), respectively. g) Representative images of CD (green) with mitochondrial (top-middle) or peroxisomal (bottom-middle) markers. The lack of overlap indicates that CD cytosolic puncta does not correspond to the tested marker and may represent an uncharacterized compartment. Scale bars represent 10 µm. h) Distribution of enzyme localization based on supporting evidence, including distinct localization, co-localization, and pathway association.

**Figure S3:**
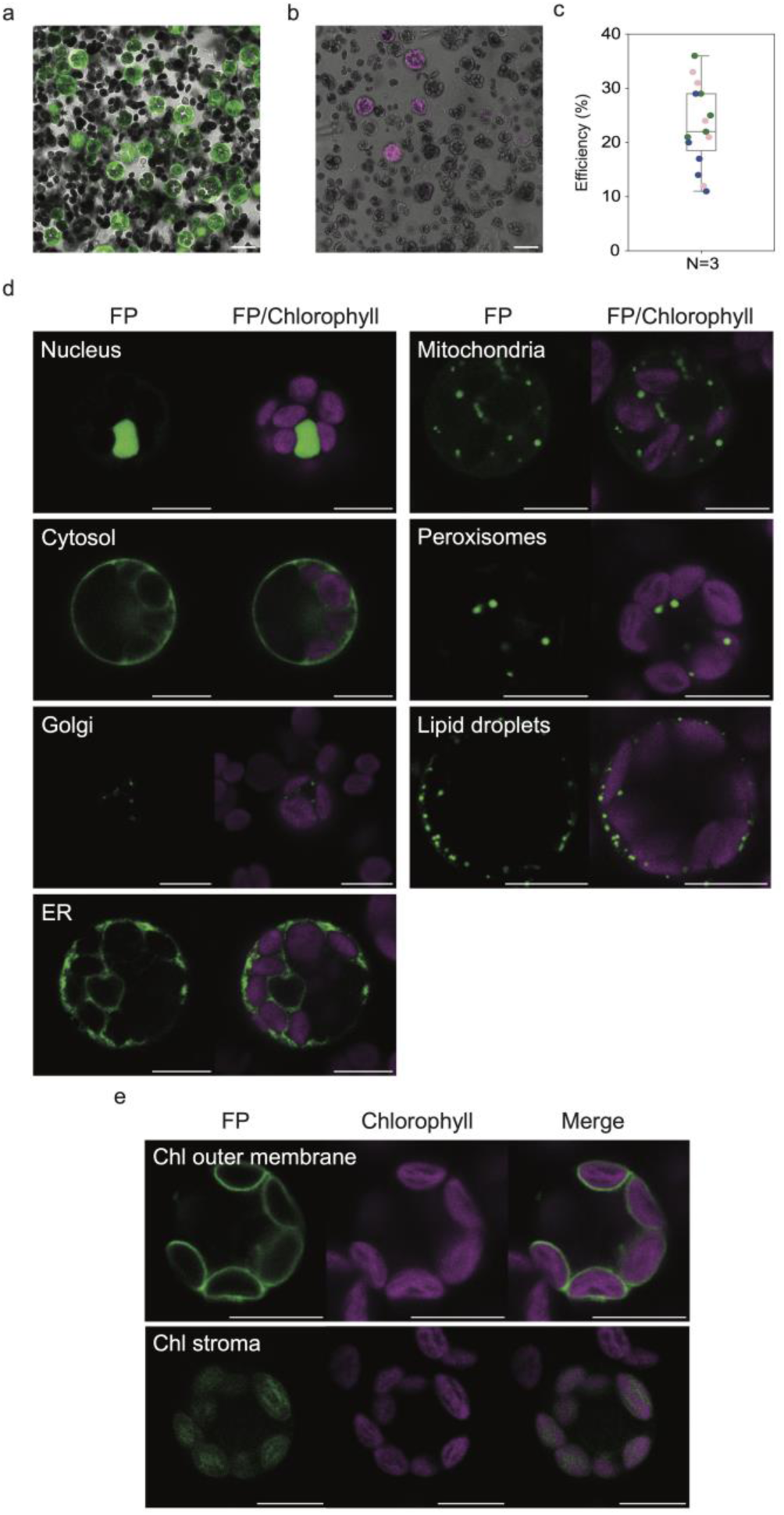
Validation of subcellular localization markers in sorghum leaf protoplasts. a) Fluorescein diacetate (FDA) staining of untransformed protoplasts. Viable protoplasts appeared as spherical, green-fluorescing cells, indicating membrane integrity. b) Representative confocal images of protoplasts transiently transformed with a mitochondrial marker (pGWTCaMV35S-Citrine-TIM21(1-50) from Osaki et al., 2017. Transformed cells expressing mitochondrial-localized Citrine are indicated by magenta merged on a bright field image. c) Quantification of transformation efficiency from five randomly captured images showed a range of 17-36% across three independent experiments. Each dot represents one image. d) FP-tagged markers for various organelles, including nucleus (pGWTCaMV35S-Citrine-NLS), cytosol (pGWTCaMV35S-Citrine), Golgi (pEZS-Sobic.004G326900-GFP as marker), endoplasmic reticulum (ER) (pEZS-𝛼AmySP-mCherry-HDEL), mitochondria (pGWTCaMV35S-TIM21(1-50)-Citrine), peroxisomes (pEZS-mCherry-PTS), and lipid droplets (Nile Red stain). Merged images display FP signals alongside chlorophyll autofluorescence where relevant. e) Fluorescent protein (FP) localization in the chloroplast outer membrane (pGWTCaMV35S-OEP7-mCherry) and stroma (pGWTCaMV35S-RBCS1a-RFP). FP signals (green) are shown separately and merged with chlorophyll autofluorescence (magenta). Scale bars represent 20 µm in panels a) and b), 10 µm in panels d) and e). Fluorescent protein signal colors have been adjusted to green for visualization.

## Supporting information

### Establishment of a protoplast transformation system in *Sorghum bicolor*

To enable large-scale fluorescent protein (FP)-based subcellular localization in *Sorghum bicolor* (sorghum), we established a method for robust protoplast isolation and transient transformation. Although FP-based transformation of protoplasts has been performed in various plant species (Gao et al., 2019; Lin et al., 2018; Trinidad et al., 2021; Wu & Hanzawa, 2018; Yang et al., 2022; Zhang et al., 2011), its implementation in sorghum remains limited (Meng et al., 2020). We optimized a protocol adapted from (Birnbaum et al., 2005; Gomez-Cano et al., 2019; Shulse et al., 2019). Protoplast viability was assessed using fluorescein diacetate (FDA) (Figure S3a), which detects esterase activity converting non-fluorescent FDA into fluorescent fluorescein in intact cells (Jones et al., 2016). Transformation efficiency was evaluated using a mitochondrion marker (TIM21 (1–50)-Citrine) driven by the CaMV35S promoter (Osaki & Kodama, 2017) (Figure S3b). Transformation efficiency ranged from 17% to 36% across three biological replicates (Figure S3c), yielding abundant protoplasts for optical screening.

To assess subcellular localization of sorghum enzymes to organelles, we assembled a panel of FP-tagged markers for major organelles, including the nucleus, cytosol, Golgi, endoplasmic reticulum (ER), mitochondria, peroxisomes, and lipid droplets. When expressed in sorghum protoplasts, subcellular localization of these markers closely resembled those observed in the plant species from which they were derived (Figure S3d). In the case of Golgi apparatus, co-localization of two enzymes with Arabidopsis orthologs that are localized to Golgi led us to select one of the enzymes as a Golgi marker (Figure S3d). Chloroplasts were readily identified by overlapping chloroplast markers and chlorophyll autofluorescence (Figure S3e). The optimized protoplast platform provides a consistent and effective approach for FP-based subcellular localization analysis in sorghum.

## Supporting Tables

**Table S1:** Construct design, fluorescent tag positioning, subcellular localization, and computational predictions for sorghum metabolic enzymes.

**Table S2:** Tissue-specific expression profiles of genes in floral, embryonic, root, shoot, and meristematic tissues.

**Table S3:** Annotation of sorghum metabolic enzymes with domain, pathway, and reaction information.

## Notes

### Competing Interest Statement

The authors have declared no competing interest.

### Summary of Updates

This revised version refocuses the manuscript as a Resource article centered on the Sorghum Metabolic Atlas and the experimentally validated subcellular localization dataset for sorghum metabolic enzymes. The revised manuscript places greater emphasis on dataset generation, experimental validation, data accessibility, and utility of the resource. The text was reorganized and revised for improved clarity and readability, including updates to the Summary, Results, Discussion, and Methods sections. Figures, figure legends, and supplemental materials were updated and reformatted accordingly.

## References

1. Ali, A.E.E. et al. (2023) ‘Comparative proteomics analysis between maize and sorghum uncovers important proteins and metabolic pathways mediating drought tolerance’, *Life (Basel*, Switzerland), 13(1), p. 170.

2. Almagro Armenteros, J.J., et al. (2019) ‘Detecting sequence signals in targeting peptides using deep learning’, Life science alliance, 2(5). Available at: 10.26508/lsa.201900429.

3. Birnbaum, K. et al. (2005) ‘Cell type-specific expression profiling in plants via cell sorting of protoplasts from fluorescent reporter lines’, Nature methods, 2(8), pp. 615–619.

4. Boccia, M. et al. (2024) ‘A scaffold protein manages the biosynthesis of steroidal defense metabolites in plants’, *Science (New York*, N.Y.), 386(6728), pp. 1366–1372.

5. Bouzroud, S. et al. (2020) ‘Down regulation and loss of auxin response factor 4 function using CRISPR/Cas9 alters plant growth, stomatal function and improves tomato tolerance to salinity and osmotic stress’, Genes, 11(3), p. 272.

6. Cutler, S.R. et al. (2000) ‘Random GFP::cDNA fusions enable visualization of subcellular structures in cells of Arabidopsis at a high frequency’, Proceedings of the National Academy of Sciences of the United States of America, 97(7), pp. 3718–3723.

7. Emenecker, R.J., Holehouse, A.S. and Strader, L.C. (2020) ‘Emerging roles for phase separation in plants’, Developmental cell, 55(1), pp. 69–83.

8. Emms, D.M. and Kelly, S. (2019) ‘OrthoFinder: phylogenetic orthology inference for comparative genomics’, Genome biology, 20(1), p. 238.

9. Gao, L. et al. (2019) ‘An efficient system composed of maize protoplast transfection and HPLC-MS for studying the biosynthesis and regulation of maize benzoxazinoids’, Plant methods, 15(1), p. 144.

10. Geraldi, A. et al. (2021) ‘Synthetic scaffold systems for increasing the efficiency of metabolic pathways in microorganisms’, Biology, 10(3), p. 216.

11. Gomez-Cano, L., Yang, F. and Grotewold, E. (2019) ‘Isolation and efficient maize protoplast transformation’, Bio-protocol, 9(16). Available at: 10.21769/bioprotoc.3346.

12. Grewal, P.S. et al. (2021) ‘Peroxisome compartmentalization of a toxic enzyme improves alkaloid production’, Nature chemical biology, 17(1), pp. 96–103.

13. Guidelli, G.V. et al. (2018) ‘pGVG: a new Gateway-compatible vector for transformation of sugarcane and other monocot crops’, Genetics and molecular biology, 41(2), pp. 450–454.

14. Hawkins, C. et al. (2021) ‘Plant Metabolic Network 15: A resource of genome-wide metabolism databases for 126 plants and algae’, Journal of integrative plant biology, 63(11), pp. 1888–1905.

15. Hooper, C., et al. (2022) ‘Subcellular Localisation database for Arabidopsis proteins version 5’. The University of Western Australia. Available at: 10.26182/8DHT-4017.

16. Hooper, C.M. et al. (2016) ‘Finding the Subcellular Location of Barley, Wheat, Rice and Maize Proteins: The Compendium of Crop Proteins with Annotated Locations (cropPAL)’, Plant & cell physiology, 57(1), p. e9.

17. Hooper, C.M. et al. (2017) ‘SUBA4: the interactive data analysis centre for Arabidopsis subcellular protein locations’, Nucleic acids research, 45(D1), pp. D1064–D1074.

18. Huh, W.-K. et al. (2003) ‘Global analysis of protein localization in budding yeast’, Nature, 425(6959), pp. 686–691.

19. Jones, K. et al. (2016) ‘Live-cell fluorescence imaging to investigate the dynamics of plant cell death during infection by the rice blast fungus Magnaporthe oryzae’, BMC plant biology, 16, p. 69.

20. Kim, W., Iizumi, T. and Nishimori, M. (2019) ‘Global Patterns of Crop Production Losses Associated with Droughts from 1983 to 2009’, Journal of Applied Meteorology and Climatology, 58(6), pp. 1233–1244.

21. Krishnakumar, V. et al. (2015) ‘A maize database resource that captures tissue-specific and subcellular-localized gene expression, via fluorescent tags and confocal imaging (Maize Cell Genomics Database)’, Plant & cell physiology, 56(1), p. e12.

22. Lacchini, E. et al. (2020) ‘CRISPR-mediated accelerated domestication of African rice landraces’, PloS one, 15(3), p. e0229782.

23. Lau, W. and Sattely, E.S. (2015) ‘Six enzymes from mayapple that complete the biosynthetic pathway to the etoposide aglycone’, *Science (New York*, N.Y.), 349(6253), pp. 1224–1228.

24. Lin, C.-S. et al. (2018) ‘Application of protoplast technology to CRISPR/Cas9 mutagenesis: from single-cell mutation detection to mutant plant regeneration’, Plant biotechnology journal, 16(7), pp. 1295–1310.

25. Mahapatra, S. et al. (2021) ‘Biofuels and their sources of production: A review on cleaner sustainable alternative against conventional fuel, in the framework of the food and energy nexus’, Energy nexus, 4(100036), p. 100036.

26. Matsumura, H. et al. (2020) ‘Hybrid Rubisco with complete replacement of rice Rubisco small subunits by sorghum counterparts confers C4 plant-like high catalytic activity’, Molecular plant, 13(11), pp. 1570–1581.

27. Meng, R. et al. (2020) ‘An efficient sorghum protoplast assay for transient gene expression and gene editing by CRISPR/Cas9’, PeerJ, 8, p. e10077.

28. Nazir, R. et al. (2022) ‘Clustered regularly interspaced short palindromic repeats (CRISPR)/CRISPR-associated genome-editing toolkit to enhance salt stress tolerance in rice and wheat’, Physiologia plantarum, 174(2), p. e13642.

29. Nett, R.S., Lau, W. and Sattely, E.S. (2020) ‘Publisher Correction: Discovery and engineering of colchicine alkaloid biosynthesis’, Nature, 584(7821), p. E35.

30. Oberortner, E. et al. (2017) ‘Streamlining the design-to-build transition with build-optimization software tools’, ACS synthetic biology, 6(3), pp. 485–496.

31. Olson, A. et al. (2014) ‘Expanding and vetting sorghum bicolor gene annotations through transcriptome and methylome sequencing’, The plant genome, 7(2), p. lantgenome2013.08.0025.

32. Osaki, Y. and Kodama, Y. (2017) ‘Particle bombardment and subcellular protein localization analysis in the aquatic plant Egeria densa’, PeerJ, 5, p. e3779.

33. Sage, R.F. (2004) ‘The evolution of C4 photosynthesis’, The new phytologist, 161(2), pp. 341–370.

34. Schindelin, J. et al. (2012) ‘Fiji: an open-source platform for biological-image analysis’, Nature methods, 9(7), pp. 676–682.

35. Schläpfer, P. et al. (2017) ‘Genome-Wide Prediction of Metabolic Enzymes, Pathways, and Gene Clusters in Plants’, Plant physiology, 173(4), pp. 2041–2059.

36. Shiferaw, B. et al. (2011) ‘Crops that feed the world 6. Past successes and future challenges to the role played by maize in global food security’, Food security, 3(3), pp. 307–327.

37. Shi, J. et al. (2017) ‘ARGOS8 variants generated by CRISPR-Cas9 improve maize grain yield under field drought stress conditions’, Plant biotechnology journal, 15(2), pp. 207–216.

38. Shulse, C.N. et al. (2019) ‘High-throughput single-cell transcriptome profiling of plant cell types’, Cell reports, 27(7), pp. 2241–2247.e4.

39. Silva, T.N. et al. (2022) ‘Progress and challenges in sorghum biotechnology, a multipurpose feedstock for the bioeconomy’, Journal of experimental botany, 73(3), pp. 646–664.

40. Tanz, S.K. et al. (2013) ‘SUBA3: a database for integrating experimentation and prediction to define the SUBcellular location of proteins in Arabidopsis’, Nucleic acids research, 41(Database issue), pp. D1185–91.

41. Thul, P.J. et al. (2017) ‘A subcellular map of the human proteome’, Science, 356(6340). Available at: 10.1126/science.aal3321.

42. Trinidad, J.L., Longkumer, T. and Kohli, A. (2021) ‘Rice protoplast isolation and transfection for transient gene expression analysis’, *Methods in molecular biology (Clifton*, N.J.), 2238, pp. 313–324.

43. Ulian, T. et al. (2020) ‘Unlocking plant resources to support food security and promote sustainable agriculture’, Plants, people, planet, 2(5), pp. 421–445.

44. Venkateswaran, K., Elangovan, M. and Sivaraj, N. (2019) ‘Origin, Domestication and Diffusion of Sorghum bicolor’, in C. Aruna et al. (eds) Breeding Sorghum for Diverse End Uses. Elsevier, pp. 15–31.

45. Wang, L. et al. (2023) ‘A chloroplast protein atlas reveals punctate structures and spatial organization of biosynthetic pathways’, Cell, 186(16), pp. 3499–3518.e14.

46. Willems, P. et al. (2024) ‘The Plant PTM Viewer 2.0: in-depth exploration of plant protein modification landscapes’, Journal of experimental botany, 75(15), pp. 4611–4624.

47. Wing, I.S., De Cian, E. and Mistry, M.N. (2021) ‘Global vulnerability of crop yields to climate change’, Journal of environmental economics and management, 109(102462), p. 102462.

48. Wu, F. and Hanzawa, Y. (2018) ‘A simple method for isolation of soybean protoplasts and application to transient gene expression analyses’, Journal of visualized experiments: JoVE [Preprint], (131). Available at: 10.3791/57258.

49. Yang, D. et al. (2022) ‘A high-efficiency PEG-Ca2+-mediated transient transformation system for broccoli protoplasts’, Frontiers in plant science, 13, p. 1081321.

50. Zhang, Y. et al. (2011) ‘A highly efficient rice green tissue protoplast system for transient gene expression and studying light/chloroplast-related processes’, Plant methods, 7(1), p. 30.

51. Zhao, C. et al. (2017) ‘Temperature increase reduces global yields of major crops in four independent estimates’, Proceedings of the National Academy of Sciences of the United States of America, 114(35), pp. 9326–9331.

52. Zhao, X. et al. (2022) ‘Colocalization analysis for cryosectioned and immunostained tissue samples with or without label retention expansion microscopy (LR-ExM) by JACoP’, Bio-protocol, 12(5), p. e4336.

